# Area-based conservation planning in Japan: protected area network effectiveness to the post-2020 Global Biodiversity Framework

**DOI:** 10.1101/2021.07.07.451416

**Authors:** Takayuki Shiono, Yasuhiro Kubota, Buntarou Kusumoto

**Affiliations:** Faculty of Science, University of the Ryukyus, Okinawa, Japan; Think Nature Inc., Okinawa, Japan; Faculty of Agriculture, Kyushu University, Japan

## Abstract

To reframe the imperfect review processes of nation-scale actions on area-based conservation through protected area (PA) networks, we first created novel infrastructure to visualize nation-level biodiversity information in Japan. We then assessed the performance of the existing PA network relative to land exploitation pressure and evaluated conservation effectiveness of PA expansion for the post-2020 Global Biodiversity Framework. The Zonation algorithm was used to spatially prioritize conservation areas to minimize biodiversity loss and the extinction risk for 8,500 Japanese vascular plant and vertebrate species under constraints of the existing PA network and land use. The spatial pattern of the identified priority areas, which were considered candidate areas for expansion of the current PA network, was influenced by land-use types according to the mask layers of non-PAs, and low-, middle-, and high-ranked PAs. The current PA network reduced the aggregate extinction risk of multiple species by 36.6%. Indeed, the percentage of built-up areas in the existing PAs was in general smaller than that in the areas surrounding PAs. Notably, high-ranked PAs fully restrained built-up pressure (0.037% per 10 years), whereas low-ranked PAs in the national park and wild-life protection areas did not (1.845% per 10 years). Conservation effects were predicted to substantially improve by expansion of high-ranked (legally strict) PAs into remote non-PAs without population/socio-economic activities, or expansion of medium-ranked PAs into agriculture forestry satoyama and urban areas. A 30% land conservation target was predicted to decrease extinction risk by 74.1% when PA expansion was implemented across remote areas, satoyama, and urban areas; moreover, PA connectivity almost doubled compared with the existing PA network. In contrast, a conventional scenario showed that placing national parks in state-owned and non-populated areas would reduce extinction risk by only 4.0%. The conservation prioritization analyses demonstrated an effectiveness of using a comprehensive conservation approach that reconciles land-sparing protection and land-sharing conservation in other effective area-based conservation measures (OECM) in satoyama and urban green spaces. Our results revealed that complementary inclusion of various PAs interventions related to their governance and land-use planning plays a critical role in effectively preventing biodiversity loss and makes it more feasible to achieve ambitious conservation targets.

## **1.** Introduction

Protected areas (PAs) are fundamental safeguards of biodiversity (Watson et al., 2014). Therefore, area-based conservation measures have been debated for several decades (Brundtland Commission, 1987; IUCN, 1994; Maxwell et al., 2020), but it remains unclear how much area or coverage and what biodiversity should be included within Pas (e.g., Chape et al., 2005). PA targets are recommended from two perspectives, biological science and policy-making, based on conservation effects (needs) and socio-economic costs (Svancara et al., 2005). Conservation scientists advocated that the 25–75% of areas should be placed within PAs to preserve biodiversity on the planet (Noss et al., 2012); alternatively, the Convention on Biological Diversity (CBD) set more feasible, conservative goals across 1992 to 2010, calling for 4–17% protection, to reduce biodiversity loss. Global Biodiversity Outlook 5 determined that the current Aichi Target 11’s 17% protection of the terrestrial surface was partly achieved in many countries by 2020; this is the most successful of all the 20 Aichi targets (CBD Secretariat, 2020a). Nevertheless, there is still rapid biodiversity loss: one million species are threatened with extinction (IPBES, 2019), which demonstrates a need for renewed, bold conservation targets to produce a biodiversity-positive outcome (Maron et al., 2018; Mace et al., 2018). Therefore, the post-2020 Global Biodiversity Framework proposes a target of 30 % land conservation (CBD Secretariat, 2020b).

Most importantly, area-based conservation targets should not include blindly placed PAs and increased spatial percentage protection motivated by political considerations or economic efficiency (Barnes et al., 2018; Magris and Pressey, 2018). Such targets should fundamentally aim for spatially prioritized PA network planning; “quality, not quantity (size), matters” (Pimm et al., 2018) means effectively capturing representativeness, ecological connectivity, and areas of importance for biodiversity patterns that are underpinned by ecological and evolutionary processes (Klein et al., 2009; Arponen, 2012; Arponen and Zupan, 2016). Therefore, biologically oriented PA expansion is a delicate mission in social equity when balancing conservation interventions and land-use policies (Moilanen and Arponen, 2011). Increasing the number of strict PAs to focus on global biodiversity, such as the ambitious suggestion of a “half-earth” initiative to protect 50% of the surface and capture approximately 85% of species (Wilson, 2016), may be infeasible in most countries with highly complicated land-use policies that involve stakeholder interaction (Büscher et al., 2016). In fact, only one-fifth of the Key Biodiversity Areas, which are defined as “sites of importance for the global persistence of biodiversity” (IUCN, 2016), are fully covered by PAs (Butchart et al., 2015), which indicates a compromised outcome behind conservation priorities.

“Other effective area-based conservation measures” (OECM) is an alternative solution to bridge the gap between socio-economic feasibility and bold conservation targets (Dudley et al., 2018; Bhola et al., 2020; Alves-Pinto et al., 2021). Notably, OECM is defined by conservation outcomes through community-led or private efforts (Jonas et al., 2014) when landscape management is shared with agricultural/forestry or indigenous traditional land use, and has less strict protection or lacks legal restriction. This conceptually may reflect the “third-of-third” conservation initiative by Hanski (2011): a third of the land area is moderately managed as multi-objective conservation landscapes, within which a third of the area is protected. For example, the rural landscape management of satoyama and urban greening in Japan provides ancillary benefits to biodiversity conservation through wildlife habitat diversification related to traditional social–ecological productions (Washitani, 2001; Kobori and Primac, 2003; Kadoya and Washitani, 2011). Therefore, a comprehensive approach that reconciles land-sparing and -sharing conservation would be a practical solution to effectively achieve more ambitious PA targets with the effectiveness (Maxwell et al., 2020). Moreover, in land-sharing OECM, securing land exploitation prevention is key for fulfilling the post-2020 Global Biodiversity Framework (Jonas et al., 2018; Donald et al., 2019); in fact, less strict Pas in satoyama and urban areas are prone to exploitation under land-use pressures (Kim et al., 2021).

Spatial conservation prioritization (SCP) supports the analytical need for PA targets in the CBD (Moilanen et al., 2014; Kullberg and Moilanen, 2014). SCP is a well-established method for identifying irreplaceable priority areas for long-term persistence of potential biodiversity and essential ecosystem services, and is based on biological and socio-economic viewpoints (Moilanen et al., 2009; Knight et al., 2011; Kukkala and Moilanen, 2013). In the SCP analysis, PA network effectiveness for biodiversity conservation is evaluated by area-dependent reduction of species extinction risks (Lehtomäki et al., 2019; Hannah et al., 2020) and spatial priority ranking across the existing PAs from intact landscape to managed (or private) landscape; this optimizes competing land uses and is computed for each area associated with stakeholder groups using their spatial layers (Moilanen et al., 2011; Honeck et al., 2020). Therefore, the SCP output facilitates outcome-based assessments of existing PA networks and guides PAs expansion by implementing top-down national-scale land-sparing and bottom-up private-led land-sharing conservation. However, the SCP-based effectiveness assessments of the PA networks have been rarely conducted in individual nations (e.g., Bicknell et al., 2017; Di Minin et al., 2017).

In this study, we measured the conservation effectiveness of the Japanese PA network: the evaluation involved the spatial design of the existing PAs, the effect of threat reduction, and the ecological outcome of PA expansion in relative to the Aichi targets and post-2020 Global Biodiversity Framework (Maxwell et al., 2020). Japan is geographically located in a global biodiversity hotspot: notably, strict PAs placed in state-owned land currently represent only 3.8% of total land areas, and most PAs (e.g., the national park) include residential, private lands, such as agriculture and forestry areas, and urban areas, which allow land exploitation; these areas differ in legal strictness and policies from those of western countries (Tanaka et al., 2019).

We first created a novel system of nation-wide biodiversity features (spatial distributions of 18,126 species, which included all vascular plants and vertebrates) and conservation cards with priority scores at the 1 km-grid resolution across the Japanese terrestrial and marine ecosystems. In this study, we used data from 8,500 terrestrial species. This information provides scientific evidence for incorporating national/regional biodiversity conservation strategies into basic plans for the conservation and sustainable use of biodiversity. Then, we investigated i) effects of exploitation pressure prevention on the existing PAs, ii) reduction of relative extinction risks by the existing PAs, iii) changes in conservation effectiveness and connectivity among the PA network by nation-level conservation actions (including the Aichi targets) over the past 20 years, and predicted iv) predicted improvements under the PA expansion scenarios related to the post-2020 target. These assessments were conducted by accounting for the existing PAs, which have different legal strictness regarding protection and different land-use types that suffer from various exploitation pressures. Finally, we propose an approach for land-sparing and -sharing conservation across landscape that includes different land-uses types, and demonstrated the importance of SCP-based PA assessment in formulating conservation targets and their measurability.

## 2. Material and methods

### 2.1. Study site

Japan mainly consists of the main islands of Japan, the Ryukyu Islands and the Bonin Islands, which are located off the eastern coast of Asia. These regions range from hemiboreal to subtropical climatic regions: the mean annual temperature ranges from – 5.3–24.2°C and the annual precipitation ranges from 650–4538 mm. This warm and wet monsoon climate contributes to development of diverse biota in the Japanese archipelago. Moreover, these islands are biogeographically located from the Holarctic to the Palaeotropical regions. The insularity during the Pliocene and Pleistocene intermittently divided one large regional biota into local communities on individual islands, with periodic connections reforming through land-bridge corridors. Such complicated geographical and climatic conditions shaped region-specific endemic biodiversity pattern through dispersal limitation and climate-related energy availability (Kubota et al., 2014, 2015, 2017). Therefore, Japan, which harbors ecologically and/or evolutionarily distinctive biota ranks among the top 35 global biodiversity hotspots (Mittermeier et al., 2011). Indeed, recent human impacts have destroyed most lowland habitats and posed threats to biodiversity hotspot conservations. Consequently, Japan serves as an ideal region where conservation biogeographers can develop spatial prioritization measures for the PA network for capturing ecological and evolutionary potential.

### 2.2. Biodiversity features

We created the Japan Biodiversity Mapping Project (J-BMP) database based on species occurrence information (n = 13,366,641 for 8,500 vascular plants and vertebrate species (Table S1). For decades, substantial knowledge of Japanese natural history has been accumulating through research activities and environmental assessments individually run by researchers, local governments, environmental assessment companies, and citizen scientists. Species occurrence, functional traits, and phylogeny data have been thoroughly compiled in J-BMP. Detailed descriptions of these data can be found on the J-BMP website (https://biodiversity-map.thinknature-japan.com/index.html).

Species distribution models were developed using Maxent version 3.4.1 (Phillips et al., 2006) with 52 environmental variables, which included climatic, soil, geological, topographical, and geographical conditions, as the predictor variables (Table. S2); these variables were previously proposed to be potentially important factors that explain biodiversity distributions in this region (Kubota et al., 2015, 2017). We binarized (1/0) the predicted suitability at the 1-km grid cell-level using the sensitivity–specificity sum maximizer threshold (Jiménez-Valverde and Lobo, 2007), and confirmed the accuracy of each model using the area under the receiver operating characteristic curve (Fig. S1). Moreover, we corrected commission errors of binarized presence/absence predictions with *ex post facto* design by validating the observed presence/absence data, such as regional species checklists, northern/southern limits of individual species distributions, and species compositions sampled at local communities (e.g., a number of vegetation samples and monitoring sites). The predicted species distributions were stacked, and species richness maps were visualized for individual taxa (e.g., plants, vertebrates, insects, stony corals, crustaceans, and shellfish etc.) on the J-BMP web site.

### 2.3. PA data

We extracted the spatial data of all PAs from Digital National Land Information (https://nlftp.mlit.go.jp/ksj/) and Natural Environmental Information GIS (http://www.biodic.go.jp/trialSystem/top_en.html) (Table S3). We classified these PA categories for biodiversity conservation into three ranks on the basis of strictness of legal protection: high = economic activities are strictly forbidden; medium = public permission is required for economic activities, and small-scale land exploitation and agriculture or forestry in private lands are allowed; and low = public permission is required for economic activities such as large-scale land exploitation and also agriculture or forestry in private lands are allowed. Subsequently, we created binary (1/0) maps of PAs for each rank at the 1-km grid cell level by defining PAs for which more than 50% protected of 100 100-m cells (100 ha) were protected. In total, approximately 20.3% of the land area was designated as PAs (high = 3.8%, medium = 6.9%, and low = 9.6%).

### 2.4. Land-use data

We compiled the data on land-use types at 1-km grid cell resolution. We first categorized individual cells into residential land-use types on the basis of the land-use subdivision mesh of the National Land Information Division, National Spatial Planning and Regional Policy Bureau, MLIT Japan (https://nlftp.mlit.go.jp/ksj/gml/datalist/KsjTmplt-L03-b.html) and human population density (https://nlftp.mlit.go.jp/ksj/gml/datalist/KsjTmplt-mesh1000h30.html). These residential land-use types included built-up areas; road and rail-way construction areas; agricultural land types, such as orchards; or forestry/grassland types, such as coastal vegetation and urban areas.

Urban areas were defined by dominance of residential land. Agriculture areas were defined as farmland areas (e.g., rice fields) in lowland areas. Satoyama areas were defined as areas for which more than 45% of the area included at least two of the following three land types based on National Actual Vegetation map (https://www.biodic.go.jp/kiso/vg/vg_kiso.html): i) agricultural land that contained natural vegetation and for which naturalness was ranked as 2 or 3, ii) secondary grass lands for which naturalness was ranked as 4 or 5, and/or iii) secondary forested lands with a predominance of *Castanopsis* or *Quercus* species and naturalness ranked as 7 or 8. The areas that were not populated, which was not applicable to urban, agricultural, and satoyama areas, were defined as remote areas, which were mostly located in mountain areas. Based on these land type classifications, we categorized all 1-km grid cells across Japan into four types of urban areas (5.8% of total land area), agricultural areas (18.0%), satoyama (45.0%), and non-populated remote mountain areas (31.2%).

### 2.5. Land exploitation prevention assessment by PA

In high-, medium-, and low-ranked PAs, we compared the composition of land-use types with their surrounding areas; the surrounding areas within a 5-km radius of existing PAs were defined as non-PAs. Moreover, to assess prevention effects of the PA network on land exploitation, we investigated built-up areas from 2006 to 2016 for each PA rank and category (e.g., natural parks, natural conservation areas, and wild-life conservation areas). To calculate naturalness score, we re-categorized land types into residential, agricultural, or forestry/grassland areas on the basis of land-use information, which included built-up areas, road and rail-way construction areas, cultivated areas, and orchards. We scored residential, agricultural, and forestry/grassland areas as 0, 1, and 2, respectively, and then counted total naturalness scores at a 100-m scale for each PA. The decrease of naturalness score from 2006 to 2016 was defined as built-up pressure: if forestry/grassland areas were replaced with residential areas, the change of naturalness score was calculated as a 100% decrease; if land-use type did not shift during this period, it was calculated as 0%. However, because we only focused on prevention of built-up pressure by PAs, we ignored increments of naturalness score (e.g., secondary succession from agricultural areas to forest/grasslands). Additionally, we evaluated the spatial distribution of solar power systems (their number and area at 1 km-grid resolution), which has recently been considered typical land exploitation in satoyama and agricultural areas with semi-natural environments (Kim et al., 2021). Spatial distribution of solar power facilities was derived from information on certification of renewable energy power generation business plans in October 2020. In this assessment, we focused on the amount of solar power facilities that generate >20 kw per 1 km2 in the existing high-, medium-, and low-ranked PA and non-PAs.

### 2.6. Spatial conservation priority analysis

We used Zonation software, which produces a balanced, complementarity-based ranking of conservation priority across the area of interest based on a set of input spatial features (Moilanen et al., 2009; Moilanen et al., 2014; Kullberg and Moilanen, 2014; Lehtomäki et al., 2019). Starting from the full landscape, Zonation iteratively ranks and removes cells that lead to the smallest aggregate loss of conservation value while accounting for total and remaining distributions of (potentially) weighted features and other relevant factors such as connectivity, costs, or habitat condition. For this analysis, we used 8,500 species distribution layers at the 1 km-grid resolution (377,589 cells), which included J-BMP-derived data from 7,346 vascular plants, 77 amphibians, 103 reptiles, 607 birds, 136 mammals, and 231 freshwater fishes. Because these groups widely cover foundation and umbrella taxa with diverse taxon-dependent biogeographic patterns (Lehtomäki et al., 2019), we assumed that they well represent the biodiversity pattern in Japan.

We conducted spatial prioritization using the continuous suitability predictions of individual species within their presence cells. In this computation, we included all species across all taxa to detect priority areas for expansion of the existing PAs using the additive benefit function (ABF) methods; this allowed achievement of cost-efficient species coverage (Moilanen et al., 2014). The ABF minimizes aggregated extinction risk and implicitly places higher priority on species-rich cells because the algorithm sums conservation values over all species in a cell (Moilanen, 2007). For individual taxa,, we weighted each species by a compound weight on the basis of the Japanese Red List category (RLC; least concern = 1, near threatened = 2, vulnerable = 4, endangered = 6, critically endangered = 8, data deficient = 2). To aggregate all species across different taxa (*j*) with different number of species (*Nj*), we normalized the aggregate weight (AGG*j*) for each taxon with *Nj* to account for the rather different numbers of species that belonged to different taxa. The weight *w* of species *i* in taxon *j* was then defined as follows: *wi, j* = (RLC*i* × AGG*j*)/*Nj*.

To identify the most cost-effective complementary additions to the current PA network, we used hierarchical mask analysis in Zonation to account for high-, medium, low-ranked PAs, remote non-PAs, satoyama areas, and urban areas. This analysis forced the highest priorities into the high-ranked PAs, followed by the medium and low-ranked PAs, finally the non-PAs of the landscape (Moilanen et al., 2011; Lehtomäki et al., 2019). Therefore, the hierarchical mask was used to implement gap analysis on the current PA network and revealed cost-efficient gap filling for future PA expansion: the top-priority, non-PAs are the most cost-effective complementary additions to the PA network (Lehtomäki et al., 2009).

### 2.7. Aggregated relative extinction risk

Using the Zonation analysis output (relationship between land loss and species’ range loss), we evaluated the relative extinction risk across multiple taxa. In the spatial prioritization of Zonation, the relative reduction (V) of distribution (R) for *i*-th species is defined as #x1D449;_"_ = #x1D445;^%^; in our analysis, the parameter *z* was set as 0.25. Therefore, species-specific extinction risk (ER*i*) relative to the proportion of PA coverage in total land areas was calculated as ER*i* =1 – R*i* . We calculated the aggregated ER over species of multi-taxon as the sum of individual species’ remaining extinction risk across all species relative to the spatial design of the PA network (Hannah et al., 2020).

### 2.8. Connectivity

PA connectivity was evaluated by the ProtConn index (Saura et al., 2017). ProtConn is defined as the percentage of a country that is covered by protected and connected lands, and includes the four fractions that account for intra- and inter-PA connectivity by moving between different PAs. ProtConn increases as area increases among connected PAs or as distance decreases between the disconnected PAs, and the maximum value of ProtConn corresponds to PA coverage (%) in total land areas. In ProtConn computation, distance thresholds were set to 10 km to represent the potential dispersial distance for species between PAs, and the probability of direct dispersal to other separated PAs was defined as 50% (Saura et al., 2018).

### 2.9. Improvement assessment: past and future PA expansion

We retrospectively and prospectively evaluated the conservation effectiveness of PA expansion retrospectively and prospectively. First, we delineated spatial distributions of PAs every 10 years during 2000–2020, and evaluated the reduction of the aggregated ER by the proportion of PA coverage and the connectivity (ProtConn) of the PA network.

Second, we simulated PA expansion following a post-2020 target (i.e., increasing PA up to 30%) based on spatial prioritization with the mask layer. To examine how the existing PA network should be expanded under the hierarchical protection system (high-, medium-, and low-ranked PAs) and heterogeneous land-use types, we built two scenarios. In the OECM scenario, we selected candidate areas from high-ranked PAs in remote areas, medium-ranked PAs in satoyama and agricultural areas, and low-ranked PAs in urban areas; these were selected based on the saturating patterns of the relative extinction curves. These expansion scenarios of new PAs across national lands in remote areas and private lands in satoyama, agricultural, and urban areas are based on the concept of combining land-sparing protection with a land-sharing approach through OECM. Alternatively, in the conventional (business-as-usual) scenario, we selected candidate expansion areas only from national land areas (national forests). In both scenarios, we evaluated their conservation outcomes as the changes in the aggregated ER and ProtConn.

## 3. Results

### 3.1. Built-up pressure prevention in PAs

The compositions of land-use type across the inside to outside of existing Pas corresponded to their legal strictness of protection (Table 1): high-ranked PAs were mostly comprised of remote areas, and medium/low-ranked PAs and the areas surrounding PAs included satoyama and agricultural areas. The percentage of built-up areas in the existing PAs was generally smaller than that in the areas surrounding Pas (Table 1): high-ranked PAs fully restrained built-up pressure, whereas low-ranked PAs in the national park and wildlife protection areas did not, especially in comparison with high- and medium-ranked PAs. In fact, the amount of solar power facilities in low-ranked PAs and/or the areas surrounding PAs was greater than that in high/middle-ranked Pas (Table 1).

**Table 1.**
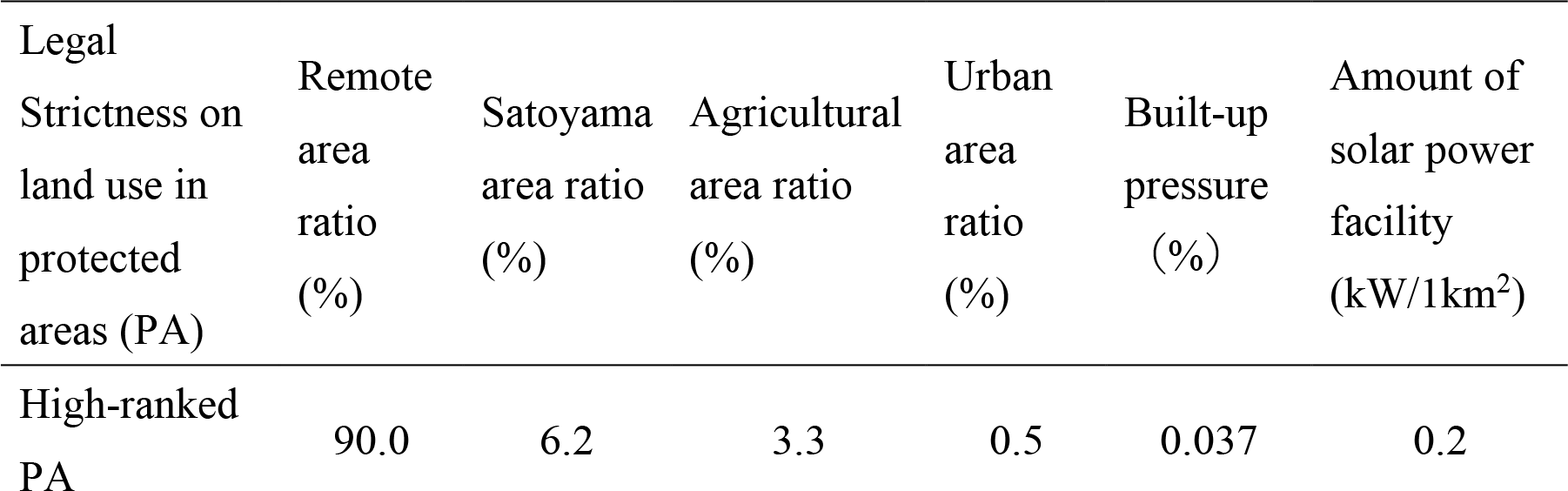
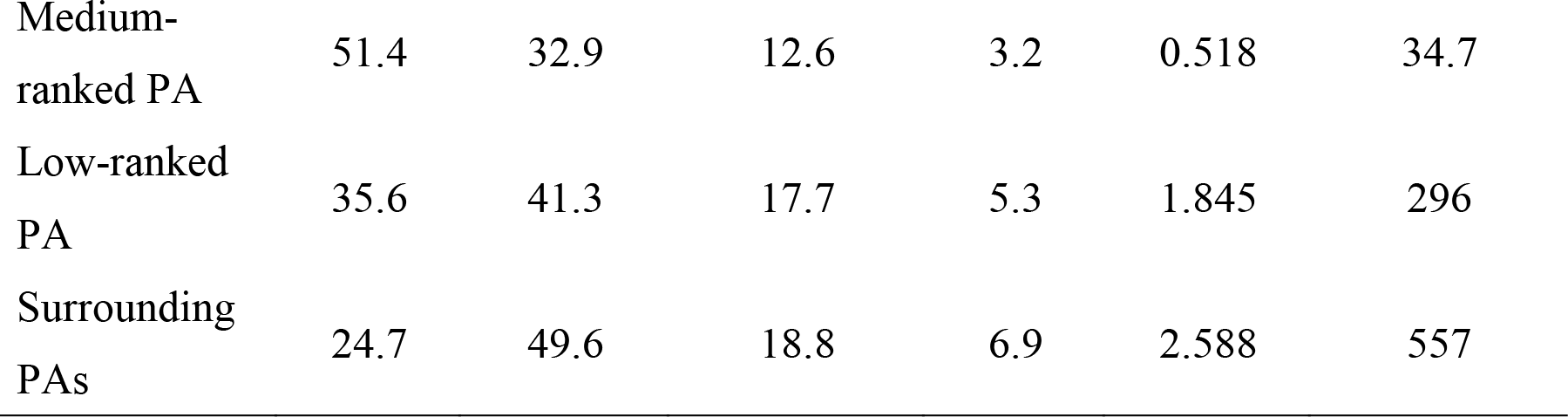
Percentage of land types on the inside and outside of the existing protected areas (PAs), built-up pressure on the existing PAs, and the amount of solar power facilities with >20kW per 1km^2^ in the existing PAs. These were assessed for high-ranked PAs (High), medium-ranked PAs (Medium), low-ranked PAs (Low), and surrounding non- PAs (Surrounding) based on land-use changes between 2006 and 2016.

| Legal<br>Strictness on<br>land use in<br>protected<br>areas (PA) | Remote<br>area<br>ratio<br>(%) | Satoyama<br>area ratio<br>(%) | Agricultural<br>area ratio<br>(%) | Urban<br>area<br>ratio<br>(%) | Built-up<br>pressure<br>(%) | Amount of<br>solar power<br>facility<br>(kW/1km<sup>2</sup>) |
| --- | --- | --- | --- | --- | --- | --- |
| High-ranked<br>PA | 90.0 | 6.2 | 3.3 | 0.5 | 0.037 | 0.2 |
| Medium-ranked PA | 51.4 | 32.9 | 12.6 | 3.2 | 0.518 | 34.7 |
| Low-ranked PA | 35.6 | 41.3 | 17.7 | 5.3 | 1.845 | 296 |
| Surrounding PAs | 24.7 | 49.6 | 18.8 | 6.9 | 2.588 | 557 |

### 3.2. Extinction risk reduction in the existing PA network

Identified conservation priority areas of terrestrial biodiversity, which included vascular plants and vertebrates were heterogeneous across Japan, especially in the middle of Honshu Island and across the Ryukyu Islands and Bonin Islands (Fig. 1): these priority areas were consistent regardless of mask layers.

**Fig. 1.**
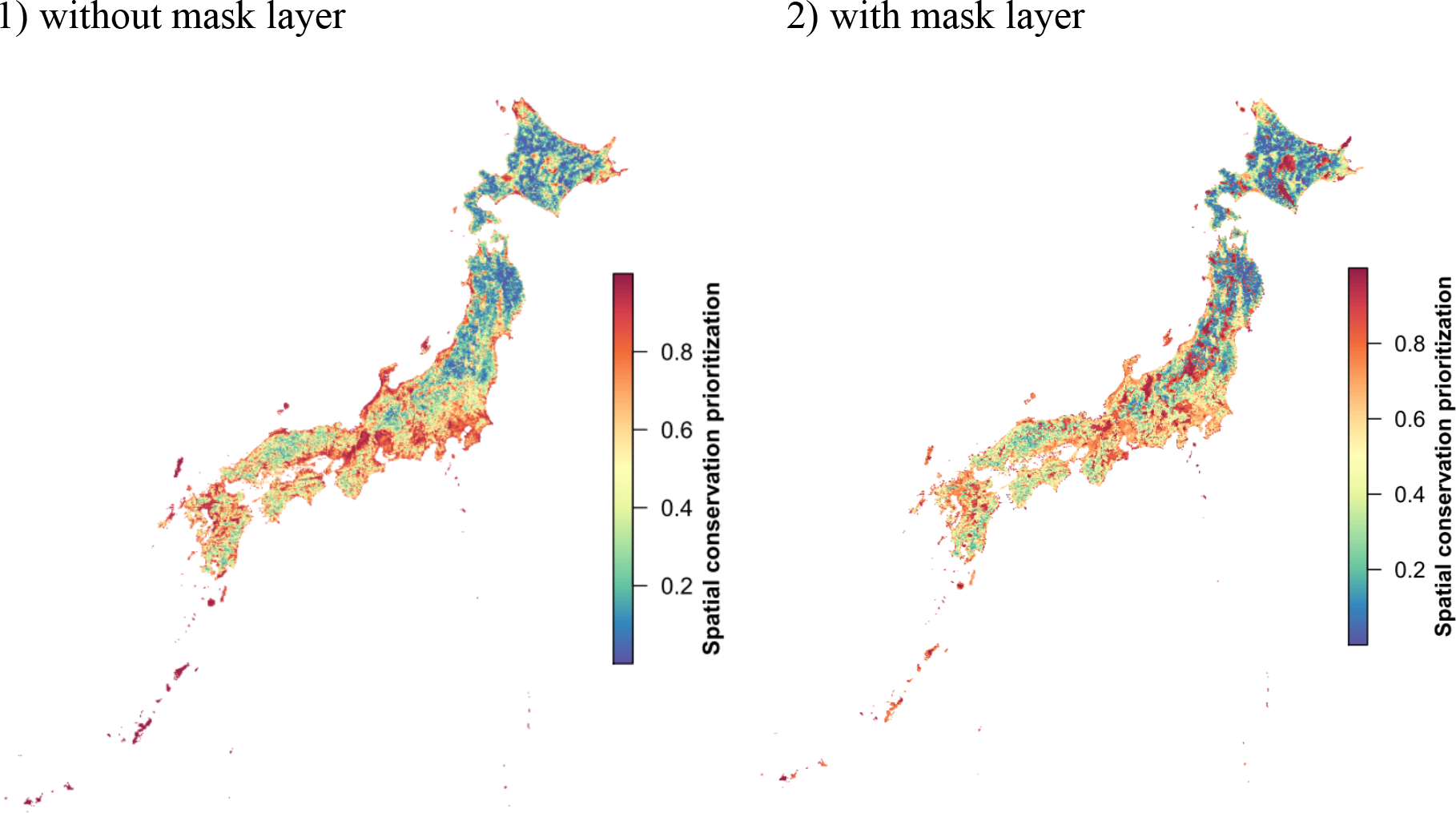
Priority areas of biodiversity conservation in Japan. Spatial conservation prioritization was based on biodiversity features across multi-taxon (8,500 species in total) and the related analysis was conducted by using the Zonation software: 1-km grid cell-level information was provided by the Japan Biodiversity Mapping Project (J-BMP; Fig. 1). The mask layer analysis accounted for high-, medium-, low-ranked protected areas (PAs), and remote non-PAs, satoyama areas, and urban areas. This analysis forces the highest priorities into the high-ranked PAs, followed by the medium- and low-ranked PAs, and finally the non-PAs of the landscape.

The existing PA network contributed to reducing the aggregate extinction risk of multiple species by 36.6% (Fig. 2): high-ranked PAs (3.73% of total land area) reduced extinction risk by 12.7%, and high- and medium-ranked PAs (10.9% of total land areas) reduced extinction risk by 25.6%. The slope of the relative extinction curve was relatively gentle within the current PAs but became steep in non-PAs. For the individual taxa (mammal, bird, reptile, amphibian, freshwater fish, and vascular plant), the aggregated relative extinction risk exponentially decreased with increasing land coverage across priority areas (Fig. S2). For all taxa, extinction risk decay rates became substantially smaller if more than 15% of land coverage was considered priority areas, and relative extinction risk was expected to reduce 30–40% (Fig. S2).

**Fig. 2.**
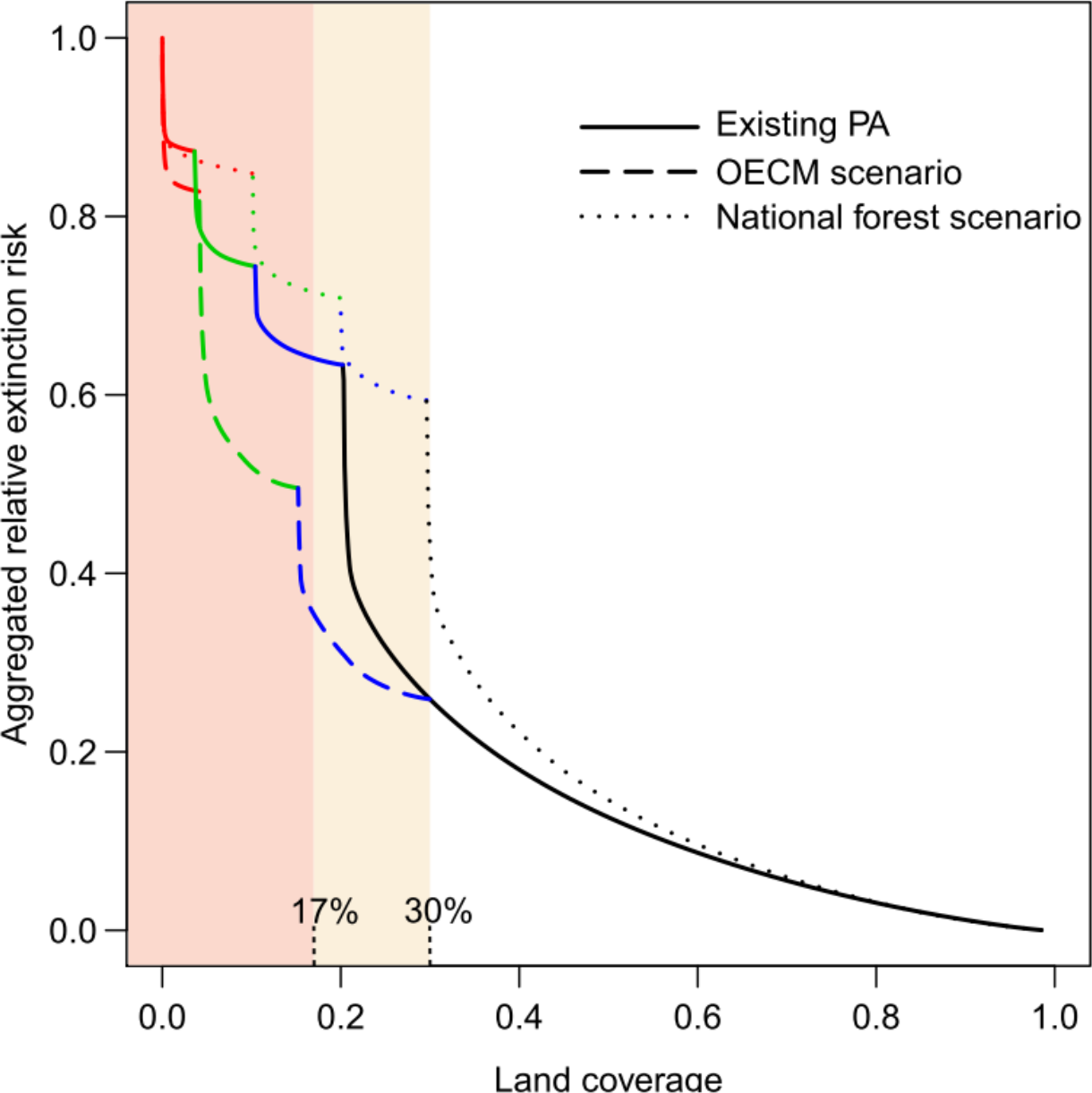
Conservation effectiveness of the current protected area (PA) network and 30% land conservation scenarios in the post-2020 Global Biodiversity Framework target that expands PAs into non-PAs. PAs effectiveness was evaluated by the reduction of aggregated relative extinction risk of all vascular plants and vertebrate species distributed in Japan. Red, green and blue curves showed conservation effectiveness of the high-, medium-, and low-ranked PAs, respectively, as increasing PA coverages across mask layers of land-use types. The solid line shows the conservation effectiveness of the current protected area. The dashed line shows the expected conservation effectiveness of a comprehensive approach that combines land-sparing protection and other effective area-based conservation measures (OECM) in satoyama and urban areas: specifically, high-ranked PAs expanded into remote state-owned areas (national forests), medium-ranked PAs into satoyama and agricultural areas, and low-ranked PAs into urban areas. The dotted line shows the expected conservation effectiveness of a conventional approach that only expanded PAs into national land areas (national forests).

### 3.3. Conservation effectiveness improvement: past and future

The total PA area increased from 74,741 to 75,892 km^2^ during 2010 to 2020. The existing PA network in 2010 reduced 35.2% of the aggregate extinction risk. The PAs expansion from 2010 to 2020, which was implemented to meet the Aichi targets, reduced the aggregate extinction risk by 1.4% and improved the PA network connectivity (Fig. 3). The spatial pattern of priority areas, which were assumed to be candidate areas to expand the current PA network, was influenced by land-use types according to the mask layers of non-PAs, and low-, middle-, and high-ranked PAs (Fig. 4). Under PA expansion up to 30% of total land area, the OECM scenario, which expands the PAs in priority areas across remote, satoyama, and urban areas, predicted 74.1% reduction in total aggregate extinction risk (Figs. 2 and 3b) and almost doubled connectivity with the current PA network (Fig. 3). Specifically, expansion of high-ranked (legally strict) PAs in remote non-PAs (0.57% of land area) without population/socio-economic activities, or expansion of medium-ranked PAs in agriculture–forestry satoyama and expansion of low-ranked PAs in urban areas (6.34% and 2.76% of land area, respectively) substantially improved conservation effectiveness. In contrast, the conventional scenario, which solely assumed PA expansion in remote, state-owned areas (e.g., national forests) reduced extinction risk by only 4.0%.

**Fig. 3.**
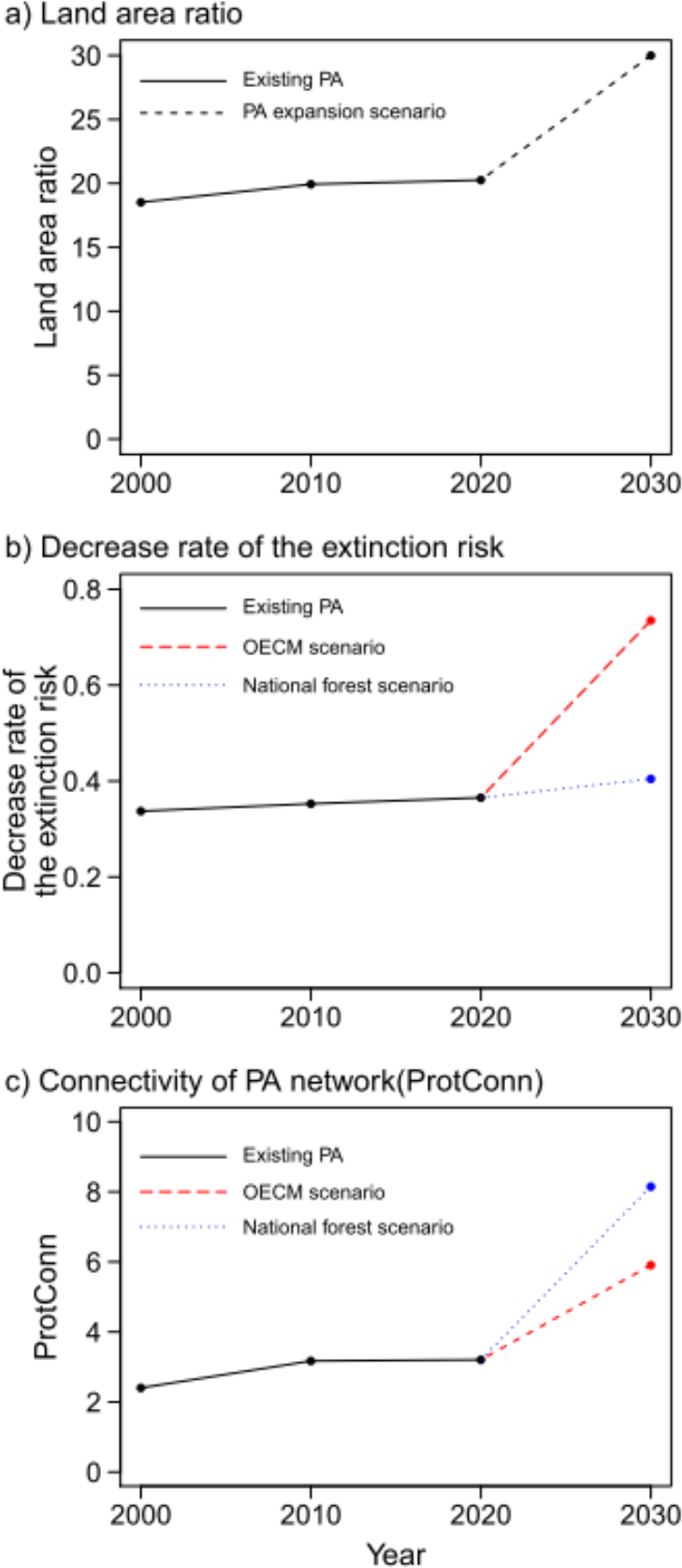
Nation-level progress of the protected area (PA) network and conservation effectiveness over the past 20 years. Conservation effectiveness was evaluated by land area ratio (%) of PAs, reduction of aggregated extinction risk, and connectivity of PAs every 10 years during 2000 to 2020. The conservation outcome of the post-2020 target (30% PAs of total land area) was predicted based on the PA expansion in priority areas across remote, satoyama, and urban areas that were identified in Fig. 2. The OECM scenario represented a comprehensive approach that combines land-sparing protection and land-sharing conservation in satoyama and urban areas, whereas national forest scenario represented a conventional approach that only expands PAs into state-owned forests.

**Fig. 4.**
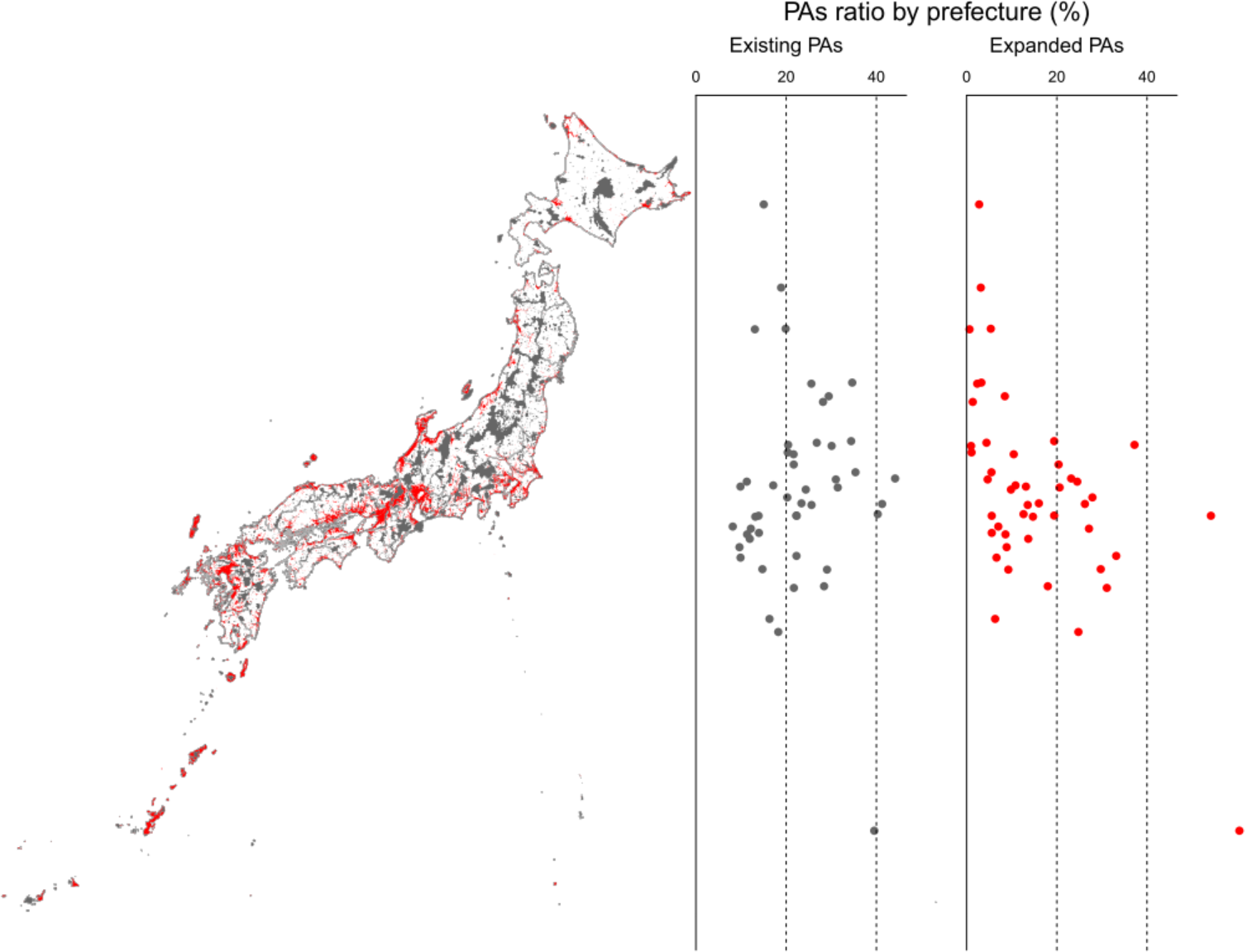
A protected area (PA) network that achieved the post-2020 Global Biodiversity Framework’s target of 30% PA coverage. Red and black areas show the expanded and existing PAs, respectively.

## **3.** Discussion

### 4.1. Conservation priority areas reduce extinction risk

Japanese nation-scale conservation priorities have been evaluated based on biological irreplaceability and the human influence index to determine threats/vulnerability (Kusumoto et al., 2017; Kubota et al., 2017; Lehtomäki et al., 2019). Additionally, the importance of area-based measures have been argued by accounting for land-use types, such as natural forests, managed forests, agricultural areas, and urban areas, which are associated with the distribution of conservation benefits and cost among local stakeholders to balance their social equity (Kubota et al., 2019). Therefore, the present SCP-based analysis measured the outcomes/effectiveness of the PA network based on the prevention of built-up pressure, the response curves of species, or aggregated extinction risk when increasing land conservation according to higher priority scores within different land-use types, which included current PAs, non-populated remote areas, satoyama (agro-forestry), agricultural areas, and urban area. Importantly, extinction risk reduction was gradual (Fig. 2), which indicated no clear thresholds such as a biologically appropriate safe limit of PA size, although the protection of 40% habitat (e.g., forest lands) to promote biodiversity persistence was empirically suggested from the relationship between habitat loss and ecological responses (Swift and Hannon, 2010; Arroyo-Rodríguez et al., 2020). Our SCP results provide scientific guidelines for nation-level actions in the framework of international (or national) conservation targets, and revealed that PAs should be maximized as much as possible to halt biodiversity loss.

### 4.2. Post-2020 PA network balancing land-sparing and -sharing conservation

The total coverage of all PAs in Japan has already achieved 20.3%, which is more than Aichi Target 11. Nonetheless, the existing Japanese PA network partially failed to prevent human impacts on priority areas: indeed, low-ranked less-strict PAs that accounted for 47.3% of total PAs were inadequate for restraining built-up pressure (Table 1), such as solar power plant deployment (Kim et al., 2021), although the current PA network reduced the aggregate extinction risk of multiple species by 36.6% (Fig. 2). This ineffectiveness of the current PA network is because high-ranked PAs are mostly located in areas with limited economic value (Kusumoto et al., 2017), as reported in nation/global-scale analyses (Venter et al., 2014; Jenkins et al., 2015). Kadoya et al. (2014) also showed that the current PA coverage is not sufficiently effective for halting extinction rate of endangered plant populations. Therefore, the conservation effectiveness of PA networks can never be achieved alone with a government-led top-down approach that focuses on nationally, owned land areas. In fact, higher ranked priority areas were substantially found privately managed lands used for agriculture or forestry and around/within urban areas that resulted in leakage from current area-based conservation planning (Ewers & Rodrigues, 2008).

Notably, the SCP outputs predicted that 30% land conservation according to priority score should reduce extinction risk by 74.1% (Fig. 2). Specifically, the expansion of high-ranked (strict) PAs in remote areas mostly without human populations and socio-economic activities (Fig. 2), and expansion of medium-rank PAs in managed satoyama and agricultural areas substantially improved conservation effectiveness by halting biodiversity loss (Fig. 2). Alternatively, the current low-ranked PAs had few restrictions on preventing land exploitation (Table 1). In fact, a number of priority areas was identified in private lands including urban areas across low-ranked PAs and non-PAs, which were distributed across Tokyo, Osaka, Kyoto, Aichi, and Fukuoka (Fig. 2), were seriously exposed to built-up pressure (Table 1). The PA expansion strategy across remote mountain, satoyama, agricultural, and urban areas contrasted with that of conventional conservation based solely on land-sparing approaches (e.g., placing national parks in state-owned areas); only expanding PAs solely in remote, non-populated areas reduced extinction risk by only a few percentage points (Fig. 2). Most national forests distributed in remote mountain areas are sufficiently captured by the existing PAs; thus, they are ineffective as candidate areas for PA expansion.

Several previous studies reported that well-managed traditional land uses are not necessarily in conflict with biodiversity conservation (e.g. Katoh et al., 2009; Jiao et al., 2019). Those studies suggested that the network of rural land areas enlarges the size of wildlife habitats and potentially plays a complementary role in area-based multi-use conservation measures. Therefore, combining private sectors’ actions in non-nationally owned areas, such as sustainable managements of rural/peri-urban landscapes, with the PA network through legal means is key to sufficiently protecting biological priority areas and implement the Satoyama initiative (http://www.env.go.jp/nature/satoyama/syuhourei/practices_en.html) (Kozar et al., 2019). Our analysis demonstrated that satoyama, agricultural, and urban areas represent substantial biodiversity, such as species in lowland waterside environments. Moreover, in a 30% area-based conservation scenario, which balances land-sparing PAs in state-owned land and OECM in satoyama, the PAs connectivity almost doubled compared with the existing PA network (Fig. 4). Therefore, a comprehensive approach that combines land-sparing protection in state-owned areas and land-sharing OECM in private land areas contributes to managing leakage and promoting resilience of the current PA network. Implementing various complementary PAs interventions related to their governance and land-use planning plays a critical role in preventing species extinction and biodiversity loss, and may makes it feasible to achieve ambitious conservation targets. Consequently, a more flexible approach across hard/soft protection measures, which include OECM (Jonas et al., 2014), community-based conservation (Brooks et al., 2012), and privately PAs (Bingham et al., 2017), is required to enhance the PA effectiveness in the post-2020 strategic goals (Bingham et al., 2017; Bhola et al., 2020).

### 4.3. Importance of nation-scale rigorous assessments on conservation effectiveness

There is a critical need for scientific assessments of the PA effectiveness in individual nations (Geldmann et al., 2013; Zafra-Calvo et al., 2017; Xu et al., 2021). All practices for reversing biodiversity decline are locally implemented by experience-based solutions and the responsibility lies within each political boundary (Arponen, 2012). Consequently, translating international commitments into individual nations’ actions is problematic because global-scale protection of biodiversity mostly depends on the details of how individual countries set spatial priorities for designing PA networks (Pimm et al., 2018). In this regard, parochialism in conservation actions should also be reconciled with transnational prioritization on the planet (Hunter and Hutchinson, 1994; Dudley, 1995; Wells et al., 2010). Nation-specific prioritization measures are more feasible and increases the local benefits of conservation, which is a positive aspect of parochialism, by accounting for region-specific constraints, such as land-use planning. However, individual nation-led collective PA expansion is relatively ineffective compared with coordinated actions promoted by cosmopolitan SCP outputs (Montesino Pouzols et al., 2014). Such an efficiency gap (negative aspects of parochialism) between globally identified priority areas and nationally prioritized/designated PAs could be collaboratively improved by SCP-based review mechanisms (Kullberg et al., 2019). Moreover, pressures on PAs are expected to intensify because of climate change and human impacts within PA boundaries or in their surrounding areas (de la Fuente et al., 2020); notably, these are side effects of land-sparing conservation or the OECM approach. Therefore, nation-specific effectiveness of land conservation is increasingly unclear or limited (Geldmann et al., 2013; Coad et al., 2019). In the post-2020 Global Biodiversity Framework, the performance measurement of PA networks should be conducted more rigorously to assist nation-specific hard/soft measures on general land-use planning, which directly supports decision-making processes of the PA expansion and management.

## Appendix. Supplementary materials

Supplementary results to this article can be found online.

## Acknowledgements

Financial support was provided by the Environment Research and Technology Development Fund (4-1501 and 4-1802) of the Ministry of the Environment, Japan. This study was also supported by the Program for Advancing Strategic International Networks to Accelerate the Circulation of Talented Researchers, the Japan Society for the Promotion of Science.

## Supplemental materials

**Table S1.** Amount of species occurrence data used to predict species distribution on the J-BMP database.

| Taxon | Number of species | Number of occurrences |
| --- | --- | --- |
| Vascular plants | 7,346 | 7,520,572 |
| Mammals | 136 | 2,049,450 |
| Birds | 607 | 3,011,635 |
| Reptiles | 103 | 71,627 |
| Amphibians | 77 | 112,109 |
| Freshwater fish | 231 | 601248 |
| Total | 8,500 | 13,366,641 |

**Table S2.** List of environment variables used to infer species distribution.

| No. | Environmental value | source |
| --- | --- | --- |
| 1 | Annual Mean Temperature | JMA's Mesh Climate Data 2000 |
| 2 | Mean Diurnal Range (Mean of monthly (max temp - min temp)) |  |
| 3 | Isothermality (* 100) |  |
| 4 | Temperature Seasonality (standard deviation *100) |  |
| 5 | Max Temperature of Warmest Month |  |
| 6 | Min Temperature of Coldest Month |  |
| 7 | Temperature Annual Range |  |
| 8 | Mean Temperature of Wettest Quarter |  |
| 9 | Mean Temperature of Driest Quarter |  |
| 10 | Mean Temperature of Warmest Quarter |  |
| 11 | Mean Temperature of Coldest Quarter |  |
| 12 | Annual Precipitation |  |
| 13 | Precipitation of Wettest Month |  |
| 14 | Precipitation of Driest Month |  |
| 15 | Precipitation Seasonality (Coefficient of Variation) |  |
| 16 | Precipitation of Wettest Quarter |  |
| 17 | Precipitation of Driest Quarter |  |
| 18 | Precipitation of Warmest Quarter |  |
| 19 | Precipitation of Coldest Quarter |  |
| 20 | Maximum snow depth |  |
| 21 | Average annual solar radiation |  |
| 22 | Total annual sunshine hours |  |
| 23 | Natural forest area | Ministry of the Environment,<br>1/50,000 Vegetation survey<br>(2nd - 5th) and 1/25,000<br>vegetation survey (6th and 7th) |
| 24 | Secondary forest area |  |
| 25 | Planted forest area |  |
| 26 | Natural grassland area |  |
| 27 | Secondary grassland area |  |
| 28 | Natural forest area within 5km |  |
| 29 | Secondary forest area within 5 km |  |
| 30 | Planted area within 5km |  |
| 31 | Natural grassland area within 5 km |  |
| 32 | Secondary grassland area within 5 km |  |
| 33 | Agricultural land area | Japanese National Land |
| 34 | Paddy area | Numerical Information, |
| 35 | City area | Landuse Subdivision Mesh |
| 36 | Wasteland area |  |
| 37 | Beach area |  |
| 38 | Agricultural land area within 5km |  |
| 39 | Paddy field area within 5km |  |
| 40 | Urban area within 5km |  |
| 41 | Wasteland area within 5km |  |
| 42 | Beach area within 5km |  |
| 43 | Land area |  |
| 44 | Water area |  |
| 45 | Elevation | Japanese National Land Numerical |
| 46 | Standard deviation of elevation | Information, Elevation and slope |
| 47 | Slope angle | angle tertiary mesh |
| 48 | Distance from the sea | Japanese National Land Numerical |
|  |  | Information, coastline |
| 49 | Geology | 1:500,000 Fundamental Land |
|  |  | Classification Survey soil map |
| 50 | Cation exchange capacity of surface soil | SoilGrid |
| 51 | Amount of organic carbon in surface soil |  |
| 52 | Surface soil pH |  |

**Table S3.**
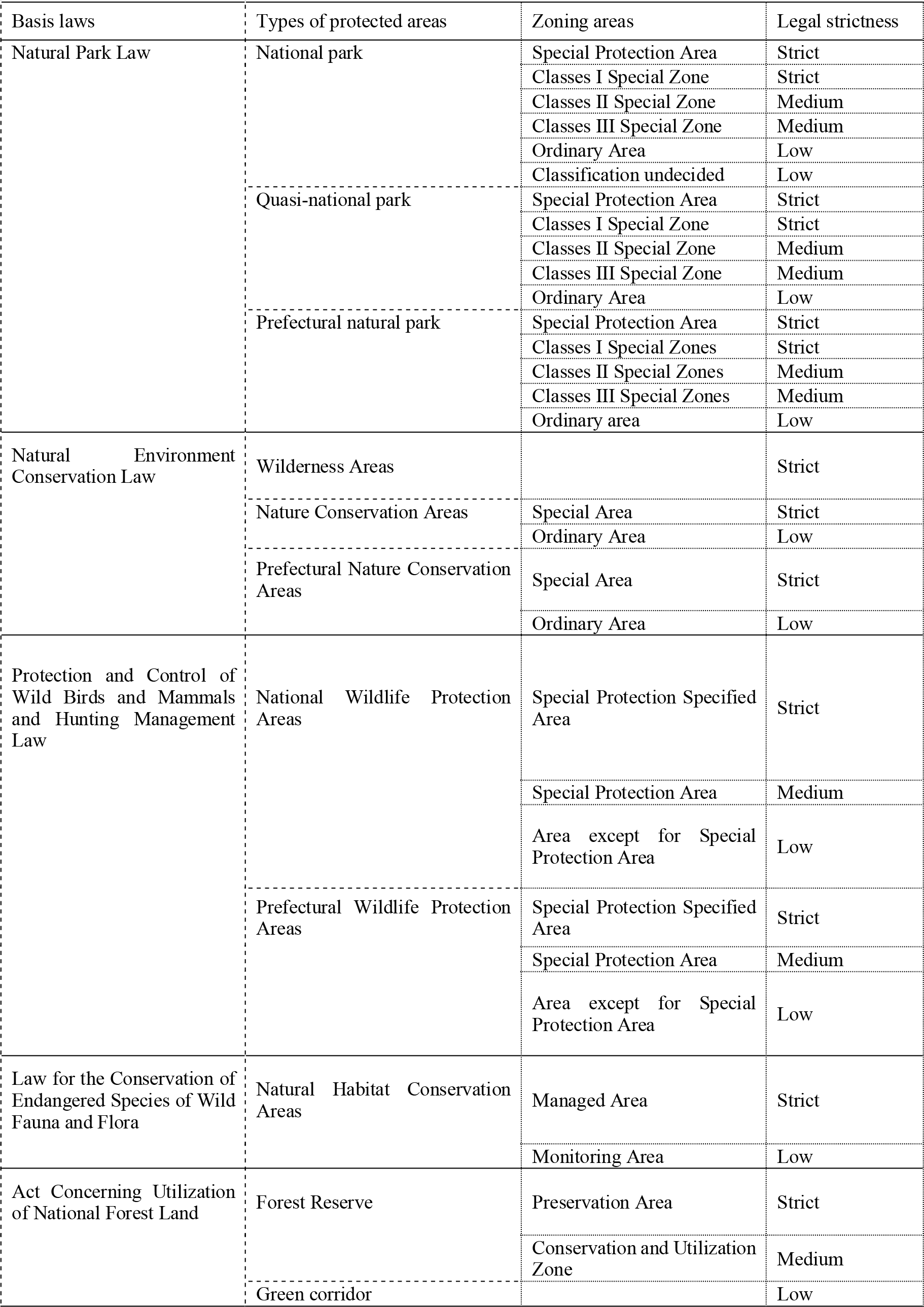
Types of Japanese protected areas and their legal strictness regarding land use.

---

**Fig S1.**
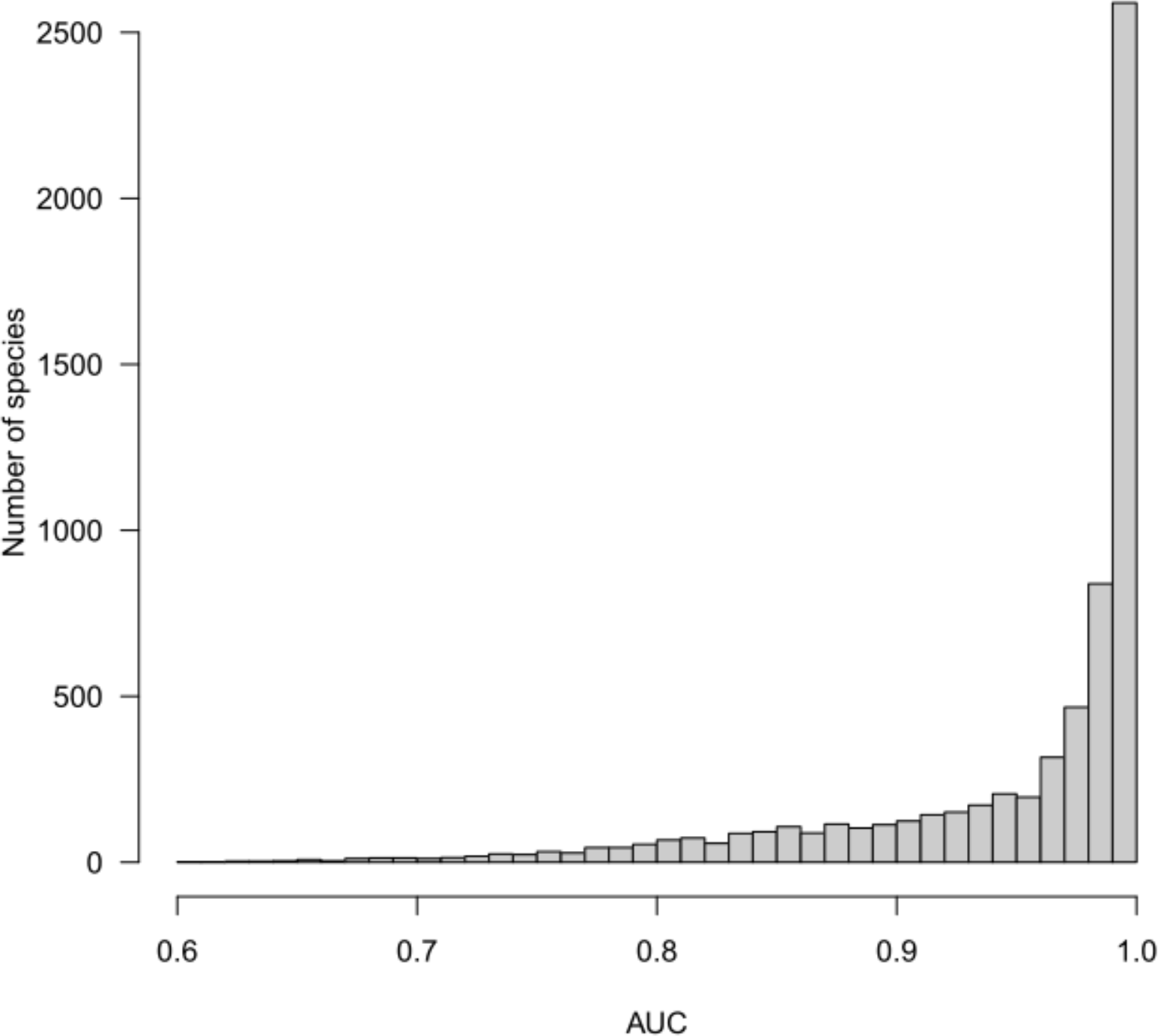
Species distribution prediction results for 8,500 species of all vascular plants and vertebrates in Japan. Mean and median values of AUC were 0.949 and 0.983, respectively.

**Fig. S2.**
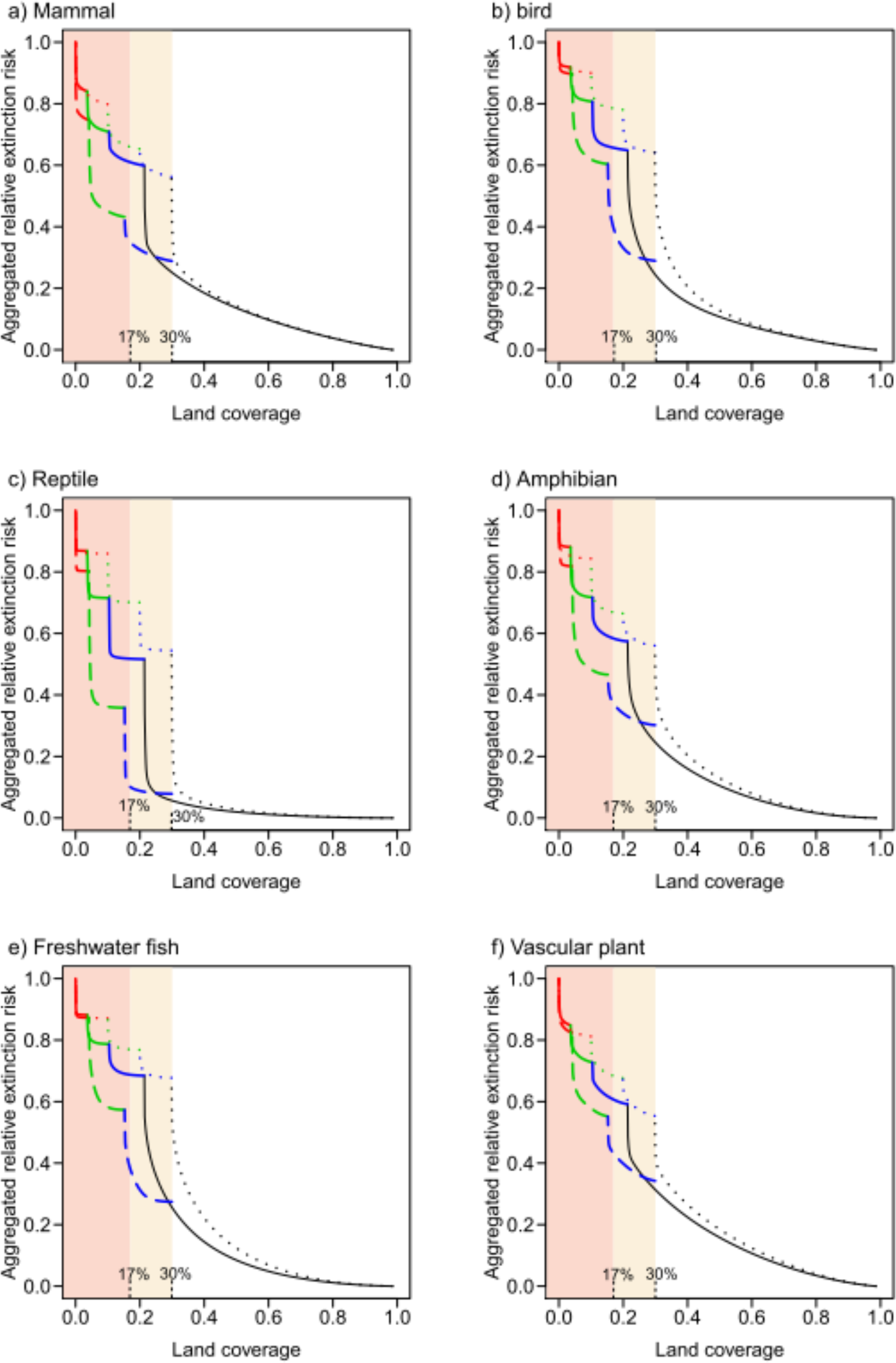
Conservation effectiveness of the current protected area (PA) network and 30% land conservation scenario in the post-2020 Global Biodiversity Framework target that expands into non-PAs. PAs effectiveness was evaluated by the reduction of aggregated relative extinction risk based on each taxon distributed in Japan. Red, green and blue curves show conservation effectiveness of the high-, medium- and low-ranked PAs, respectively, with increasing PA coverage across mask layers of land-use types. The solid line shows the conservation effectiveness of the current PA network. The dashed line shows the expected conservation effectiveness of a comprehensive approach that combines land-sparing protection and other effective area-based conservation measures (OECM) in satoyama and urban areas: specifically, high-ranked PAs expand into remote state-owned areas (national forests), medium-ranked PAs expand into satoyama and agricultural areas, and low-ranked PAs expand into urban areas. The dotted line shows the expected conservation effectiveness of a conventional approach that only expands PAs into national land areas (national forests).

**Fig. S3.**
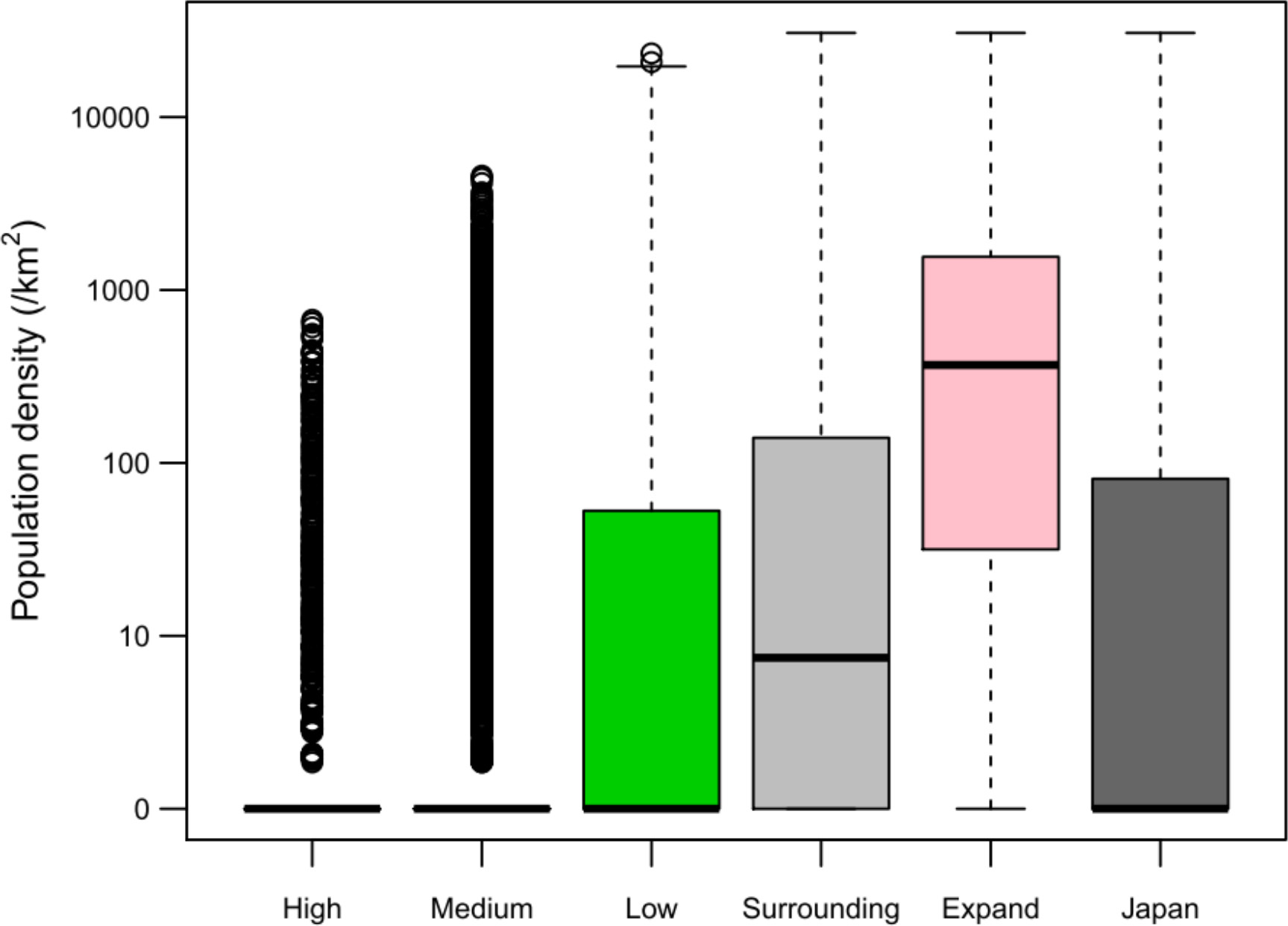
Population density on the inside and outside of existing protected areas (PAs). Population density (/km^2^) was evaluated in high-ranked PA (High), medium-ranked PA (Medium), low-ranked PA (Low), surrounding non-PAs (Surrounding), expanded PAs under the post-2020 Global Biodiversity Framework target (Expand), and all over Japan based on land types (Japan).

